# Polyploid genome assembly of *Cardamine chenopodiifolia*

**DOI:** 10.1101/2024.01.24.576990

**Authors:** Aurélia Emonet, Mohamed Awad, Nikita Tikhomirov, Maria Vasilarou, Miguel Pérez-Antón, Xiangchao Gan, Polina Novikova, Angela Hay

**Affiliations:** Max Planck Institute for Plant Breeding Research, Carl-von-Linné-Weg 10, 50829 Köln, Germany; Plant Science Research Laboratory (LRSV), UMR5546 CNRS/University of Toulouse 3, 24 chemin de Borde Rouge, 31320 Auzeville Tolosane, France; State Key Laboratory for Crop Genetics and Germplasm Enhancement, Bioinformatics Center, Academy for Advanced Interdisciplinary Studies, Nanjing Agricultural University, 210095, Nanjing, China

**Keywords:** *Cardamine chenopodiifolia*, polyploidy, amphicarpy, explosive seed dispersal, chromosome-based long-read assembly

## Abstract

**Background:** *Cardamine chenopodiifolia* is an amphicarpic plant that develops two fruit morphs, one above and the other below ground. Above-ground fruit disperse their seeds by explosive coiling of the fruit valves, while below-ground fruit are non-explosive. Amphicarpy is a rare trait that is associated with polyploidy in *C. chenopodiifolia*. Studies into the development and evolution of this trait are currently limited by the absence of genomic data for *C. chenopodiifolia*.

**Results:** We produced a chromosome-scale assembly of the octoploid *C. chenopodiifolia* genome using high-fidelity long read sequencing with the Pacific Biosciences platform. We successfully assembled 32 chromosomes and two organelle genomes with a total length of 597.2 Mbp and an N50 of 18.8 kbp (estimated genome size from flow cytometry: 626 Mbp). We assessed the quality of this assembly using genome-wide chromosome conformation capture (Omni-C) and BUSCO analysis (97.1% genome completeness). Additionally, we conducted synteny analysis to infer that *C. chenopodiifolia* likely originated via allo-rather than auto-polyploidy and phased one of the four sub-genomes.

**Conclusions:** This study provides a draft genome assembly for *C. chenopodiifolia*, which is a polyploid, amphicarpic species within the Brassicaceae family. This genome offers a valuable resource to investigate the under-studied trait of amphicarpy and the origin of new traits by allopolyploidy.

## Introduction

*Cardamine chenopodiifolia* Pers. (NCBI:txid 3101730) is an annual flowering plant that belongs to the Brassicaceae family and is native to, and widespread in, South America (Cabrera, 1967; Cheplick, 1983; Gorczyński, 1930; Persoon, 1807). *C. chenopodiifolia* is amphicarpic, meaning it bears fruit both above and below ground, with two very distinctive modes of seed dispersal (Fig. 1). Exploding seed pods are produced above ground and disperse their many, small seeds by explosive coiling of the fruit valves. Another type of seed pod develops below ground. The few, large seeds produced by each of these fruits are dispersed underground. The flowers on the main shoot of *C. chenopodiifolia* are positively geotropic and immediately grow towards the soil. These reduced flowers self-pollinate while growing through the soil and develop fruit that set seed underground. In contrast, the axillary shoots of *C. chenopodiifolia* grow away from gravity and produce flowers that typically self-pollinate, but can also be cross pollinated, and develop explosive fruit. The unique biology of *C. chenopodiifolia* makes it an ideal species to study the development and evolution of the unusual trait of amphicarpy.

**Figure 1:**
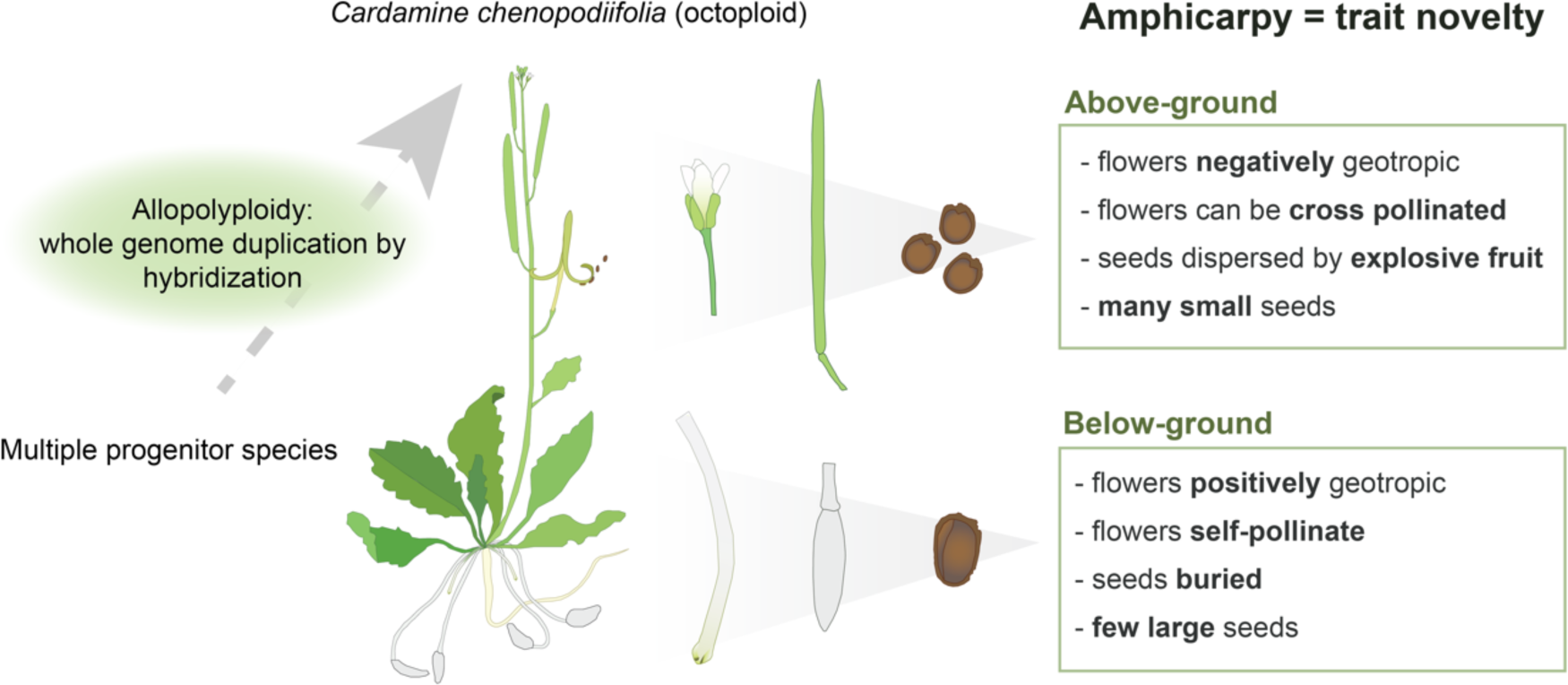
*C. chenopodiifolia* is an amphicarpic species. Amphicarpic plants bear aerial and subterranean fruit with distinct dispersal strategies and reproductive characters.

Cardamine is one of the largest genera in the Brassicaceae with more than 200 species (Al-Shehbaz, 1988; Marhold et al., 2018), among which 58% are described as polyploids (Kučera et al., 2005). Based on chromosome counts, *C. chenopodiifolia* has been described almost 100 years ago as an octoploid (Manton, 1932). Another octoploid species in this genus, *C. occulta*, was recently sequenced as a model for ruderal weeds (Li et al., 2023). The diploid species *C. hirsuta* is commonly used as an experimental system for comparative studies with its relative *Arabidopsis thaliana* (Baumgarten et al., 2023; Hay and Tsiantis, 2016; Hay et al., 2014; Hofhuis et al., 2016; Monniaux et al., 2018; Vlad et al., 2014). The reference genomes and vast array of genetic tools in these two model plants make it attractive to develop emerging model species within this phylogenetic neighborhood. This provides another motivation to assemble the octoploid genome of *C. chenopodiifolia* as a valuable tool for comparative studies and polyploidy research.

Here, we report the first whole-genome assembly for *C. chenopodiifolia* using single-molecule real-time sequencing technology from Pacific Biosciences (PacBio HiFi), and Omni-C technology. We show that the *C. chenopodiifolia* genome is octoploid and comprised of four sub-genomes, each with eight chromosomes. One sub-genome was more distinct than the others, allowing us to phase this set of chromosomes by gene tree topology analysis.

## Material and methods

### Growth conditions

*Cardamine chenopodiifolia* seeds (Ipen: XX-0-MJG-19—35600) were obtained from the Botanic Garden of the Johannes Gutenberg University, Mainz, Germany, and self-pollinated for five generations to ensure homozygosity. Aerial seeds were germinated in long-day conditions on ½ Murashige and Skoog (MS) plates after 7 days of stratification, then 1-week-old seedlings were transferred to soil and grown in a walk-in chamber (16h light, 20 °C; 8h dark, 18°C; 65% humidity). *Cardamine hirsuta*, herbarium specimen voucher Hay 1 (OXF), was cultivated on soil in long-day conditions (LD; days: 20 °C, 16 h; nights: 18 °C, 8 h) after stratification on soil at 4°C in the dark for 7 days.

### Chromosome spreads

Mitotic chromosome spreads were performed as previously described (Cromer et al., 2019) with minor modifications. Inflorescences were immediately fixed in fresh 3:1 Clarke’s fixative (3 vol. absolute ethanol: 1 vol. acetic acid). Fixative was refreshed 3 times. After fixation, inflorescences were dissected under a binocular, and white closed buds were collected (no pollen present). Samples were washed twice 2 min in deionized water, and twice 2 min in 10 mM trisodium-citrate buffer (pH4.5, adjusted with HCl). Samples were digested for 1 hour 45 min at 37°C (digestionmix: 0.3% (w/v) Pectolyase Y-23 (MP Biomedicals), 0.3% (w/v) Driselase (Sigma), 0.3% (w/v) Cellulase Onozuka R10 (Duchefa), 0.1% sodium azide in 10 mM tri-sodium-citrate buffer). Three to six buds were transferred on a clean slide in a drop of water and dilacerated with a thin needle until it formed a suspension. 10 µl of 60% acetic acid was then delicately incorporated into the suspension with a hooked needle, and then the slide was heated on a hot block at 45°C for 1 min while slowly stirring. Another 10 µl of 60% acetic acid was added as the drop started to evaporate and stirred for a supplementary minute. To mount the slide, ice-cold fresh fixative solution was pipetted as a boundary around the droplet and allowed to invade the slide. Then, a jet of ice-cold fixative was applied twice directly onto the center of the circle. After the removal of excess fixative by tilting, the slide was dried at room temperature. Once dried, 8 µl of DAPI solution (2 µg/ml in antifade mounting medium Citifluor AF1, Agar Scientific) was added onto a coverslip and mounted on the slide. Imaging was performed using a Zeiss Axio Imager Z2 microscope and Zen Blue software. Images were acquired with a Plan-Apochromat 100×/1.40 Oil M27 objective, Optovar 1.25× Tubelens. The excitation and detection windows for DAPI were set as follows: emission, 335–385 nm, detection, 420-470 nm.

### Flow cytometry

Flow cytometry was performed according to a modified protocol from Doležel et al. (1992). To release nuclei, newly expanded leaves of *C. hirsuta* and *C. chenopodiifolia* were chopped with a sharp razor blade on a petri dish containing 300 µL of Galbraith’s buffer (45 mM MgCl_2_, 20 mM MOPS, 30 mM sodium citrate, 0.1 % (v/v) Triton X-100) (Galbraith et al., 1983) and including 50 mg/L RNAase. Nuclei suspensions were passed through 50 µm CellTrics® filters and stained with propidium iodide (PI) at a final concentration of 50 mg/L for 1 h on ice in darkness. Stained nuclei of *C. hirsuta* were analyzed separately and in combination with *C. chenopodiifolia* in a CytoFLEX (Beckmann Coulter) platform using the excitation and emission parameters for PI. 10 000 events were recorded for each sample and gating was employed to exclude doublets and debris. Gating, analysis, and plotting were performed using the manufacturer’s software (CytEXPERT).

Estimated genome size was calculated with the following formula where *GS = genome size, 2C = mean peak position* (PI-area):

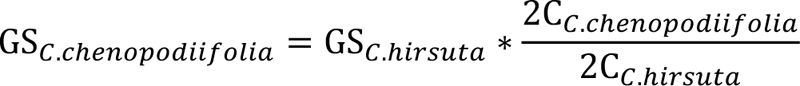

### DNA extraction, library construction, and sequencing

For genome sequencing, high-molecular-weight (HMW) DNA was isolated from 2 g fresh, shock-frozen seedlings (liquid nitrogen) with the NucleoBond HMW DNA kit (Macherey Nagel, Düren, Germany) and DNA quality was assessed by capillary electrophoresis (Agilent FEMTOpulse). For PacBio library preparation, the HMW DNA was fragmented with g-tubes (Covaris) to get 20 kbp fragments and then a library was prepared according to the recommendations of the SMRTbell Express Template Prep Kit 2.0 (Pacific Biosciences). Next, a size selection was applied to enrich for >=10 kbp fragments (BluePippin, Sage Sciences), followed by long-read sequencing on four SMRT cells on a Sequel II device with Sequel II Binding kit 2.0, Sequel II SMRT 8M cells, and Sequel II Sequencing Plate 2.0 chemistry for 30 hours and a final concentration of 110 pmolar on plate. Sequencing was performed at the Max Planck Genome-centre Cologne. In parallel, a chromatin-capture library (Omni-C, Dovetail) was prepared according to recommendations from the vendor, followed by 2 x 150 paired-end sequencing on an Illumina NextSeq 2000 device at Max Planck Genome-centre Cologne resulting in 71599541 reads.

### Assembly of C. chenopodiifolia genome

We used GALA (gap-free long-read assembly tool, version 1.0.0) for *de novo* assembly of the *C. chenopodiifolia* genome (Awad and Gan, 2023). Since GALA uses preliminary assemblies to cluster long reads into multiple groups for chromosome-by-chromosome data analyses, three draft assemblies were constructed using 85× coverage PacBio Hifi reads using HiCanu v2.1 (Koren et al., 2017), Flye v2.4 (Kolmogorov et al., 2019) and Hifiasm 0.5-dirty-r247 (Cheng et al., 2021). Assembly was conducted using default parameters and an expected genome size of 600 Mbp.

The straightforward application of GALA generated 35 scaffolding groups. Among them, two scaffolding groups were assembled into single contigs with telomeric motifs at one end, indicating each group represented a chromosome arm. We thus merged these two groups into a single scaffolding group and performed single-chromosome assembly using the LGAM module of GALA. The assembly of GALA was gap-free and complete, containing 32 pseudomolecules and 2 organelle chromosomes.

We then polished the GALA assembly to enhance the assembly correctness. The HiFi raw reads were mapped to the GALA assembly using minimap2 v. 2.17-r941 and the command ‘minimap2 -x asm20’ (Li, 2018). Then, we used an in-house genome polisher to enhance the correctness of the assembled genome.

Minimap2 was used with the command-line option “-ax asm5” to map the final assembly of *C. chenopodiifolia* against the published reference genome of *C. hirsuta* (Gan et al., 2016) to phase the 32 assembled chromosomes into four representative sub-genomes.

### Assembly quality validation

Assembly contiguity was assessed with a Python script (see code availability). Assembly completeness was assessed by Benchmarking Universal Single-Copy Orthologs (BUSCO v.3.0.0) (Seppey et al., 2019) with the dataset Embryophyta_odb9. To assess correctness, we used ‘minimap2 -x asm20’ to map the HiFi reads to the final assembly. Then, we collected the mapping statistics from samtools-stats (Danecek et al., 2021; Li et al., 2009). Finally, we called the variants and collected the variant calling statistics using BCFtools (Li, 2011). To evaluate collapsing, we collected the depth information using samtools depth and marked all the regions with depth = average depth*2 as collapsed regions. Finally, we used Python to plot the average value of window size 5000 bp. The Omni-C data were analysed with Juicer v1.6 (Durand et al., 2016b) and the contact map was visualized in Juicebox (Durand et al., 2016a).

### Alignment to *C. chenopodiifolia* transcriptome

*C. chenopodiifolia* long and short read transcripts (obtained from https://www.ebi.ac.uk/ena/browser/view/PRJEB69676) were aligned to the *C. chenopodiifolia* assembly using HISAT2 v2.1.0 (Kim et al., 2019).

### Synteny analysis

We assessed the large-scale similarity of chromosomes in the assembly using the wgd v2.0.22 pipeline (Chen and Zwaenepoel, 2023) with a draft annotation obtained from the Helixer web interface (https://www.plabipd.de/helixer_main.html). We made the required CDS sequence file (as well as the proteome file for further analyses with OrthoFinder) using the AGAT v1.0.0 GFF parser (agat_sp_extract_sequences.pl). We clustered these sequences into gene families using the wgd dmd command, calculated the synonymous substitution rate between family members with wgd ksd, and visualized these Ks values on a self-synteny map made by i-AdHoRe v3.0 (Proost et al., 2012) using wgd syn. We visualized the results using R v4.2.2 with the tidyverse v2.0.0 data manipulation suite (Wickham et al., 2019). Some repetitive tasks were sped up with GNU parallel (Tange, 2011). All software was run under default settings.

### Paralog divergence analysis

We used the output of wgd ksd to visualize the distribution of paralog divergence in the *C. chenopodiifolia* genome, filtering out pairs of genes found on the same chromosome or having an alignment coverage (alignment length divided by the larger gene length) ≤0.5 in order to reduce the noise from small-scale gene duplications and mis-grouped genes. This set of gene pairs was split into two parts, one with pairs involving the “most diverged” sub-genome (as inferred by gene tree analysis, see Results) and another with the remaining gene pairs.

### Gene tree topology analysis

Taking the *Crucihimalaya himalaica* proteome (Zhang et al., 2019) as an outgroup, we inferred the rooted orthogroup gene trees for *Cardamine chenopodiifolia* and *Crucihimalaya himalaica* proteomes using OrthoFinder v2.5.5 (Emms and Kelly, 2019). We used these trees to find the most frequently observed relationships between the homeologs using Newick Utilities v1.1.0 (Junier and Zdobnov, 2010) and gotree v0.4.5 (Lemoine and Gascuel, 2021). For this purpose, the tips of each tree were generalized to the chromosome where the gene is located using nw_rename, and redundant tips were then collapsed using nw_condense. Among the condensed trees, for each of the eight sets of four homeologous chromosomes (as found by synteny analysis) we counted trees matching each of 15 possible rooted four-leaf topologies using nw_match.

### Data availability

The PacBio and Omni-C raw read data and the genome assemblies generated by GALA in this study have been deposited at European Nucleotide Archive (ENA) PRJEB71776.

### Code availability

The script for assessing assembly contiguity is available at: https://github.com/mawad89/assembly_stats. The source code of GALA is available from GitHub at https://github.com/ganlab/GALA under the MIT license.

## Results

### *C. chenopodiifolia* genome is octoploid

To verify the ploidy of the *C. chenopodiifolia* plants used for sequencing, we performed chromosome spreads using mitotic cells of flower buds and counted 64 chromosomes (Fig. 2A). This suggests that *C. chenopodiifolia* has 8 sets of 8 chromosomes, typical of the ancestral crucifer karyotype (Lysak et al., 2006). We then estimated genome size by flow cytometry, comparing the nuclei DNA contents of *C. chenopodiifolia* with *C. hirsuta* as standard reference. *C. hirsuta* has a genome size of 255 Mbp (Gan et al., 2016) and its nuclei were separated into five peaks representing a DNA content of 2C, 4C, 8C, 16C, and 32C (Fig. 2D-E). Pooling nuclei from both species for flow cytometry allowed us to compare their relative DNA content. We identified 3 peaks belonging to *C. chenopodiifolia*, representing DNA content of 2C’, 4C’ and 8C’ (Fig. 2B-C). The first 2C’ peak of *C. chenopodiifolia* had a mean position (PI-area = 159474.8) slightly lower than the 8C peak of *C. hirsuta*, suggesting that its genome is octoploid (Fig. 2B, D). We estimated the genome size of *C. chenopodiifolia* to be 626 Mbp (225 Mbp × (159474.8 / 57315.6), see Methods).

**Figure 2:**
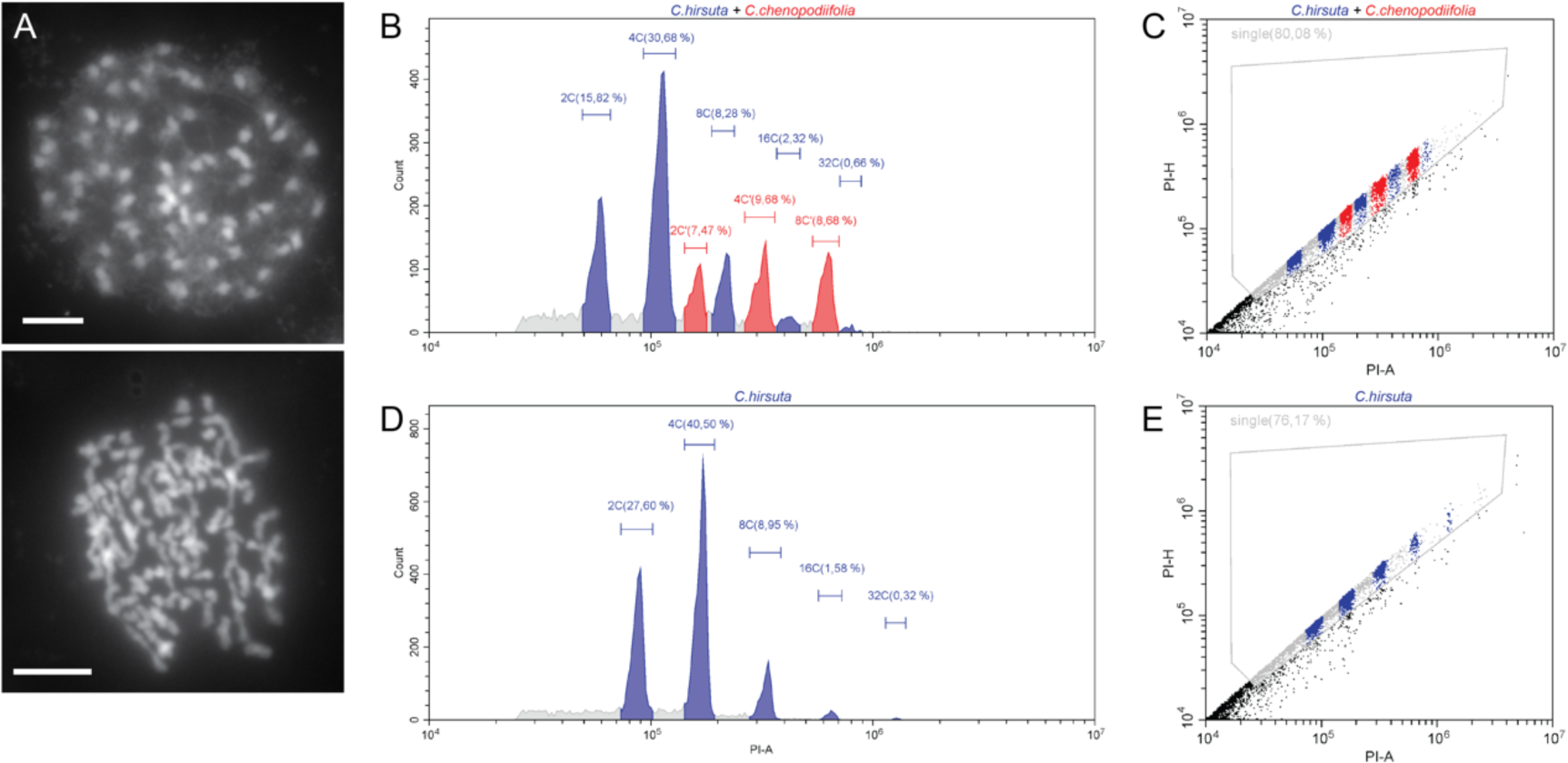
*C. chenopodiifolia* is octoploid and has 64 chromosomes. **(A)** Metaphase spreads in an aerial *C. chenopodiifolia* flower cell stained with DAPI. Two examples are shown. **(B-E)** Flow cytometry analysis of a mixture of *C. hirsuta* and *C. chenopodiifolia* (B-C) or *C. hirsuta* alone (D-E). (B, D) The X-axis indicates the area of the signal from propidium iodide (PI-A) on a logarithmic scale. The Y-axis indicates the number of nuclei recorded (Count). Segments were drawn manually to cover each peak. Blue: *C. hirsuta*. Red: *C. chenopodiifolia*. The percentage of events corresponding to each peak over the total events is indicated. XC indicates the endopolyploidy level (e.g.: 2C - nuclei in a diplophasic state). (C, E) Single nuclei events were discriminated and by assessing propidium iodide fluorescence area (PI-A) versus height (PI-H).

### *C. chenopodiifolia* chromosome-level genome assembly

To ensure homozygosity, *C. chenopodiifolia* plants were self-pollinated for 5 generations by single seed descent using aerial seeds, before sequencing. High molecular weight DNA was extracted from seedlings and sequenced using the PacBio Sequel II platform. We generated 103 Gb of raw DNA sequence (corresponding to 85× coverage) comprising 2978449 reads with a mean read length of 17.28 kbp (Table 1, Fig.3A). Long reads were pre-assembled with HiCanu, Flye and Hifiasm softwares. We used these draft assemblies with GALA (Awad and Gan, 2023) to obtain a final assembly by chromosome-to-chromosome analysis without the need of Omni-C data. This resulted in an almost complete chromosome-level genome of 597 Mbp with N50 value of 18.8 kbp (Table 2). We obtained 32 chromosomes, one mitochondrial genome, and one plastid genome. Only one gap was left in the centromeric region of chromosome 9 (Table 2).

**Table 1:** PacBio HiFi dataset parameters.

| **HiFi reads** |  |
| --- | --- |
| **Total bases (Gb)** | 103 |
| **Total reads** | 2978449 |
| **Read N50 (kbp)** | 17.19 |
| **Read mean (kbp)** | 17.28 |
| **Read L50** | 1225263 |
| **Coverage** | 85 x |

**Table 2:** Statistical data for the *Cardamine chenopodiifolia* genome.

| **Parameters** | **Values** |
| --- | --- |
| Genome size (bp) | 597'266'279 |
| Chromosomes | 32 + 2 organelles |
| N50 (kbp) | 18.80 |
| L50 | 15 |
| N75 (kbp) | 16.99 |
| L75 | 23 |
| Longest Chr (kbp) | 24.07 |
| Gaps | 1 |
| Ploidy | Octoploid |

**Figure 3:**
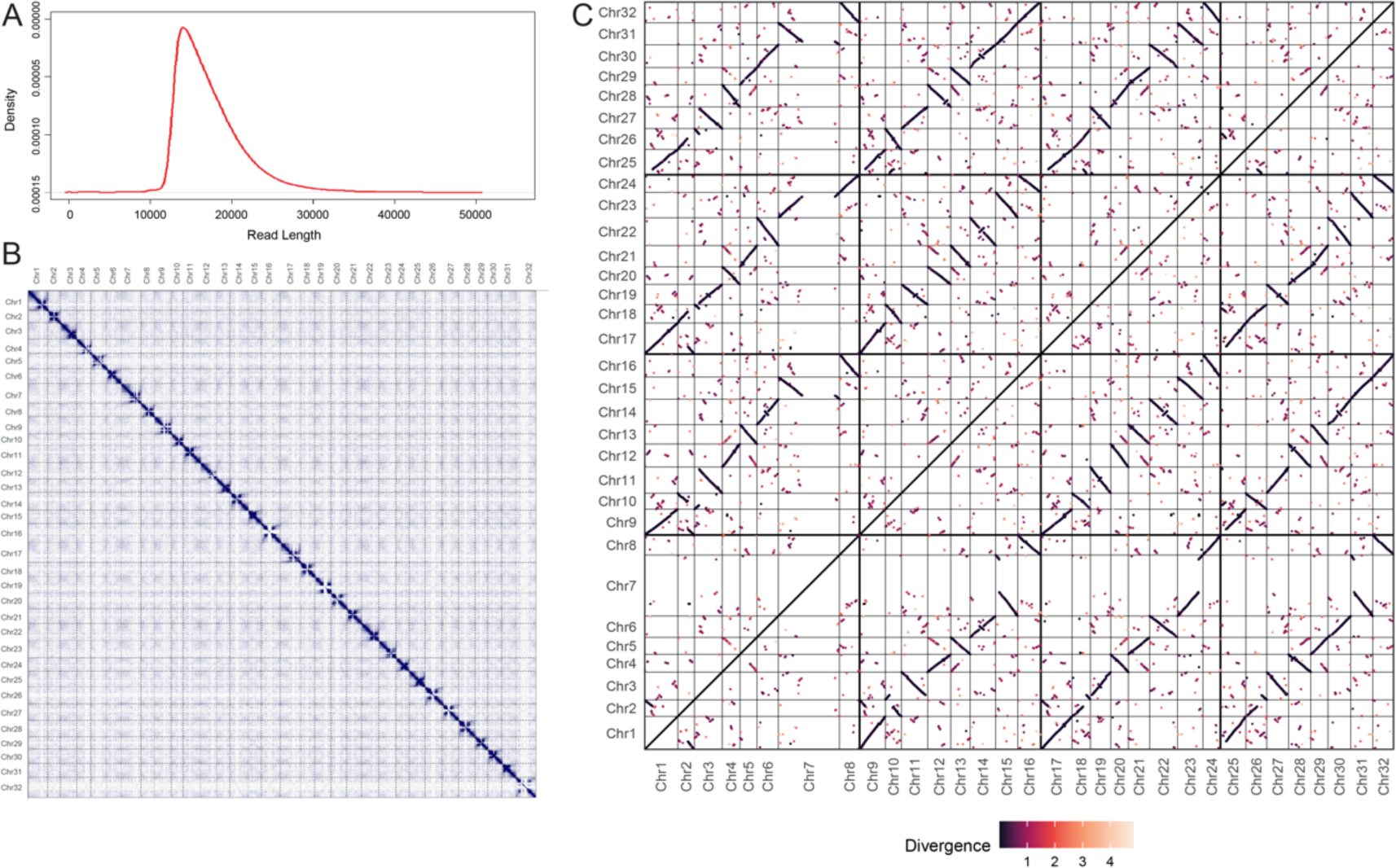
Assembly of 32 gap-free chromosomes. **(A)** HiFi Read density. **(B)** Omni-C contact matrix showing no mis-assembly. **(C)** Self-collinearity dotplot of the *C. chenopodiifolia* assembly. Each dot represents a pair of highly similar genes, the color reflecting their divergence (expressed in a number of synonymous substitutions per synonymous site, or Ks). Dark lines at Ks < 1 reflect the genome multiplication that happened in the Cardamine ancestors of *C. chenopodiifolia*. Thick lines delimit four groups of homeologous chromosomes. Note the large private inversions in some homeologs, e.g. in Chr28 as compared to Chr4, Chr12, and Chr20.

### Assessment of genome quality

Hi-C technology allows chromosomal structure to be linked directly to genomic sequence and can, therefore, be used to achieve chromosome-scale scaffolding. Since our initial assembly had already resolved the 32 chromosomes of *C. chenopodiifolia*, we used our Omni-C data to assess its quality: 71599541 raw Omni-C reads were mapped to the genome assembly using Juicer to generate a contact matrix. The contact maps results showed 32 unambiguous chromosomes with no obvious mis-assemblies (Fig. 2B).

We also evaluated the completeness of the genome assembly using Benchmarking Universal Single-Copy Ortholog (BUSCO). 97.1% of the 1,440 conserved core Embryophyta genes were identified as complete (28.7% as single-copy genes and 68.3% as duplicated genes). Additionally, 0.6% of the genes were recognized as fragmented (Table 3).

**Table 3:** Benchmarking Universal Single-Copy Orthologs (BUSCO)

|  | Number of genes |
| --- | --- |
| **Complete BUSCOs** | 1397 (97.1%) |
| **Single-copy BUSCOs** | 414 (28.7%) |
| **Duplicated BUSCOs** | 983 (68.3%) |
| **Fragmented BUSCOs** | 8 (0.6%) |
| **Missing BUSCOs** | 35 (1.7%) |

When we mapped PacBio HiFi reads to the genome assembly with minimap2, we found the overall mapping rate was 99.98%, indicating that most of the sequencing data was represented (Table 4). We plotted the distribution of coverage depth for the whole genome (Figs. S1, S2). The coverage was regular along all chromosomes except for some regions with lower coverage on chromosomes 4, 5, 14, and 32, and with higher coverage on chromosomes 9, 12, and 28, indicating extended and collapsed sequences, respectively.

To test the correctness of our assembly, we used BCFtools for variant calling (Table 4). A low number of variants is usually a sign of a correctly assembled genome. We obtained only 20800 variants, most of them being insertions or deletions, suggesting that our genome assembly is generally correct and is highly homozygous.

**Table 4:**
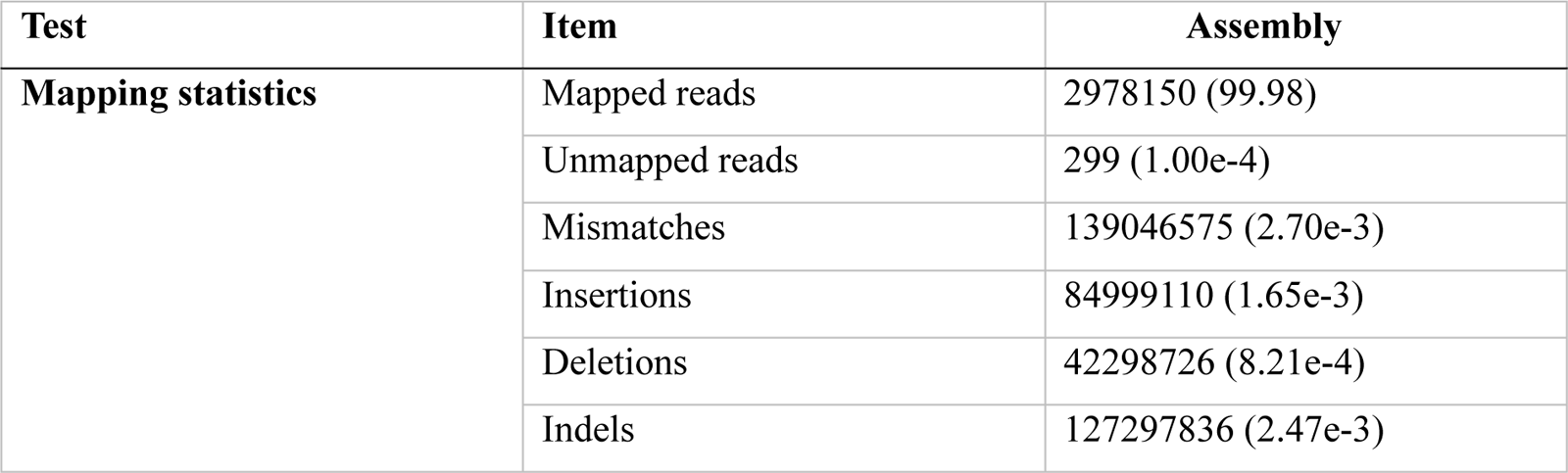
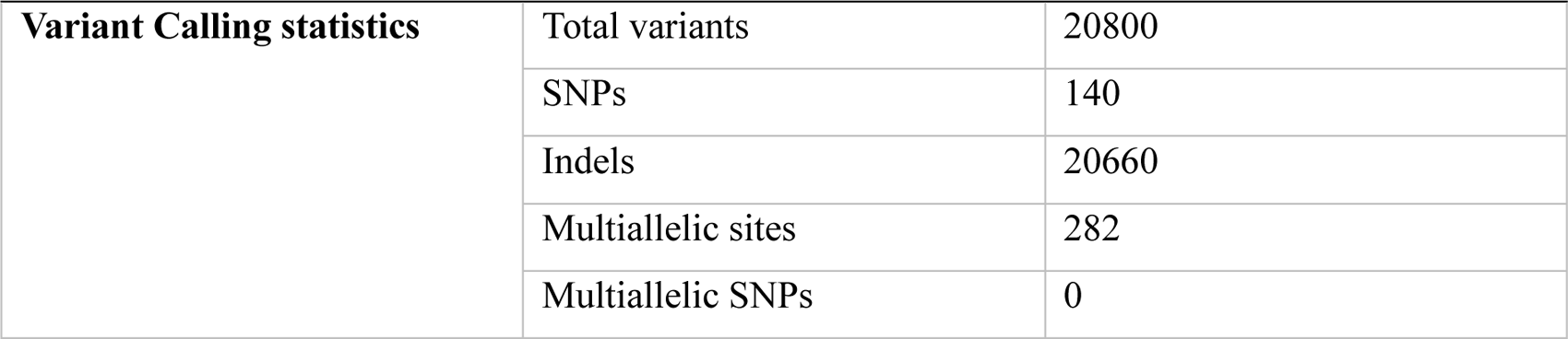
HiFi read mapping and variant calling.

| Test | Item | Assembly |
| --- | --- | --- |
| **Mapping statistics** | Mapped reads | 2978150 (99.98) |
|  | Unmapped reads | 299 (1.00e-4) |
|  | Mismatches | 139046575 (2.70e-3) |
|  | Insertions | 84999110 (1.65e-3) |
|  | Deletions | 42298726 (8.21e-4) |
|  | Indels | 127297836 (2.47e-3) |
| **Variant Calling statistics** | Total variants | 20800 |
|  | SNPs | 140 |
|  | Indels | 20660 |
|  | Multiallelic sites | 282 |
|  | Multiallelic SNPs | 0 |

Additionally, we used HISAT2 to align RNA-seq data from *C. chenopodiifolia* to the genome assembly. The transcriptome data comprised of Illumina short reads and PacBio Iso-Seq long reads (https://www.ebi.ac.uk/ena/browser/view/PRJEB69676). Short reads aligned from 91.64 to 94.04% while long reads aligned from 93.08% to 97.56%.

### *C. chenopodiifolia* is likely to be an allooctoploid

In an attempt to group the chromosomes from our genome assembly into sub-genomes, we first compared *C. chenopodiifolia* chromosomes to *C. hirsuta* chromosomes by mapping our final assembly to the published reference genome of *C. hirsuta* (Gan et al., 2016). We obtained eight sets of four chromosomes, with each set mapping to one of the eight *C. hirsuta* chromosomes (Table 5). This suggests that the *C. chenopodiifolia* genome is composed of four sub-genomes, but does not elucidate their relationships.

Polyploids have more than two sets of chromosomes and are typically classified as auto- or allopolyploids, depending on their evolutionary history. Autopolyploids arise by whole genome duplication within a single species. In contrast to this, allopolyploids are the result of a hybridization event between different species (Stebbins, 1947; Van de Peer et al., 2017). To investigate ploidy and distinguish between these two scenarios in *C. chenopodiifolia*, we first assessed the large-scale similarity of chromosomes by synteny analysis (Fig. 3C). We observed profound structural variation among all homeologs, which shows that different homeologs are unlikely to recombine and thus supports allopolyploidy. For example, there were instances where one of the four homeologs had a private big inversion, as seen in Chr28 as compared to Chr4, Chr12 and Chr20 (Fig. 3C).

We then aimed to find the most frequently observed relationships between homeologous chromosomes. Therefore, we generated rooted orthogroup gene trees for *C. chenopodiifolia* and *Crucihimalaya himalaica* (outgroup) proteomes (Zhang et al., 2019). We found that in each of the eight sets of homeologs, one chromosome was preferentially resolved as sister to the three other chromosomes, suggesting that it belongs to the most diverged sub-genome (Fig.4A). Three tree topologies matching that condition were almost equally frequent in most of the homeologous groups. This means that although the four sub-genomes are dissimilar, as suggested by the synteny analysis, their relationships are poorly resolved beyond the early-diverged sub-genome. Re-running OrthoFinder with the more sensitive IQ-TREE v2.2.2.1 tree inference software (Minh et al., 2020) did not improve the resolution.

**Figure 4:**
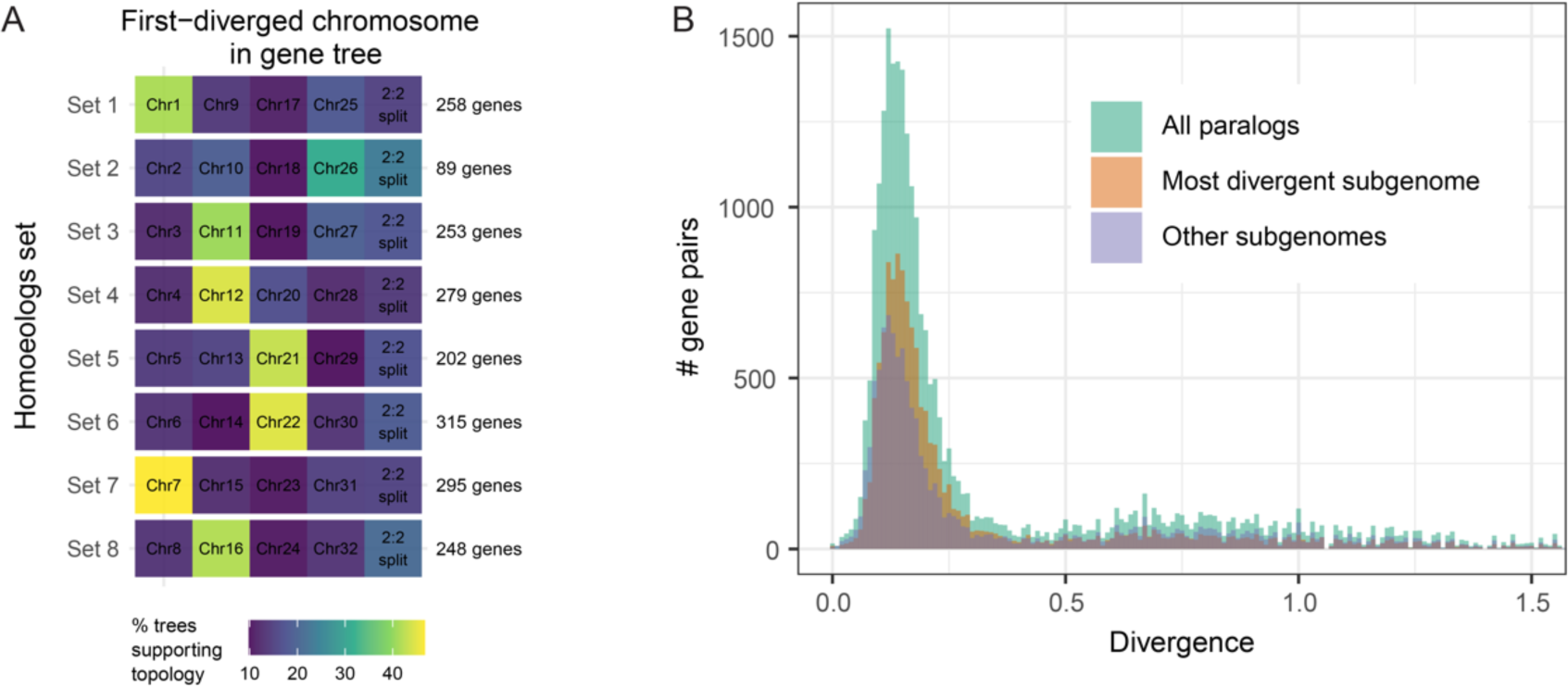
*C. chenopodiifolia* is allooctoploid. **(A)** Relative frequency of five tree topology categories (columns) among eight sets of homeologous chromosomes (rows). In each row, there is one dominant topology category that suggests the existence of one most divergent chromosome. **(B)** Distribution of paralog divergences (expressed in number of synonymous substitutions per synonymous site, or Ks) in the *C. chenopodiifolia* genome. The dataset from the entire paranome (green) is split into pairs including the most divergent sub-genome (orange) and the combinations of genes from other sub-genomes (purple).

The divergence spectrum of paralogous genes appeared to form a single peak (Fig. 4B), contrary to the expectation of distinct sub-genome divergence events suggested from the synteny analysis above. However, the pairs involving the “most diverged” sub-genome, as found by tree topology analysis, formed a peak with the maximum slightly shifted towards higher divergence, corroborating the lower similarity of this sub-genome to the others, and suggesting that the one visible peak might be formed by a few smaller overlapping peaks produced by comparisons of similarly diverged sub-genomes (Fig. 4B).

In conclusion, the *C. chenopodiifolia* genome is composed of four different sub-genomes. We could confidently phase one sub-genome and show it comprised chromosomes 1, 26, 11, 12, 21, 22, 7 and 16, with a one-to-one correspondence to chromosomes 1 to 8 of *C. hirsuta*, respectively (Table 5).

**Table 5:** Correlation of the *C. chenopodiifolia* chromosomes to the closest *C. hirsuta* chromosome. One sub-genome was phased by tree topologies and paralog divergence analysis (Fig. 4).

|  | *C. chenopodiifolia* chromosomes |  |
| --- | --- | --- |
| *C. hirsuta* chromosomes | Phased sub-genome | Unphased sub-genomes |
| **Chr1** | Chr1 | Chr9, Chr17, Chr25 |
| **Chr2** | Chr26 | Chr2, Chr10, Chr18 |
| **Chr3** | Chr11 | Chr3, Chr19, Chr27 |
| **Chr4** | Chr12 | Chr4, Chr20, Chr28 |
| **Chr5** | Chr21 | Chr5, Chr13, Chr29 |
| **Chr6** | Chr22 | Chr6, Chr14, Chr30 |
| **Chr7** | Chr7 | Chr15, Chr23, Chr31 |
| **Chr8** | Chr16 | Chr8, Chr24, Chr32 |

## Discussion

Under-studied species are an important resource in light of climate change and biodiversity challenges. Investigation of trait diversity requires the characterization of unconventional model organisms and the establishment of genomic tools in these species. The Brassicaceae family is rich in species characterized by very diverse traits and different ecological adaptations (Nikolov et al., 2019). Moreover, this family contains the model species *Arabidopsis thaliana* and many other species with sequenced genomes, thus providing rich genomic resources for comparative studies. This motivated us to develop the polyploid species *Cardamine chenopodiifolia* as an emerging model organism in the Brassicaceae to study the unusual trait of amphicarpy.

We provide here a chromosome-level genome assembly for *C. chenopodiifolia*. Using the PacBio platform, we generated a 597.2 Mbp octoploid genome assembly composed of 32 chromosomes (2n = 64), with an N50 length of 18.8 kbp. Both the estimated ploidy and genome size from this assembly agreed well with our results from cytology and flow cytometry.

We used this *C. chenopodiifolia* genome assembly to investigate its polyploid origin. Octoploids are usually expected to arise from the whole genome duplication of a tetraploid, or by the hybridization of two different auto- or allotetraploid species. These scenarios are considered more likely due to the challenges inherent to successful meiosis in a newly formed polyploid (Bomblies and Madlung, 2014). Our synteny analysis revealed many rearrangements between *C. chenopodiifolia* homeologous chromosomes, which is an unexpected scenario for an autopolyploid with random chromosome pairing. In addition, there was no clear pattern of pairwise similarity as expected from a merger of two divergent autotetraploid genomes. Therefore, we think that *C. chenopodiifolia* might have four dissimilar sub-genomes, resulting, for example, from the merger of two allotetraploids with non-overlapping parental species.

Polyploids make up more than half of the large number of species in Cardamine (Kučera et al., 2005; Marhold et al., 2018), and hybridization between species has been previously reported for a number of these polyploids. For example, the allotriploid *C. insueta* and the allotetraploid *C. flexuosa*, are hybrids between *C. amara* and *C. rivularis*, and *C. amara* and *C. hirsuta*, respectively (Akiyama et al., 2021; Sun et al., 2020). Differences between the sub-genomes of *C. chenopodiifolia*, coupled with low heterozygosity in this selfing plant, might have facilitated the chromosome-level haploid assembly that we produced with minimal involvement of Hi-C scaffolding. As suggested by gene trees and the paralog divergence spectra, one of the sub-genomes is sister to the three other sub-genomes, whose relationships could not be resolved confidently. Future work will focus on phasing the remaining three sub-genomes and providing an annotation for the assembly. This is likely to help identify possible progenitors of *C. chenopodiifolia*.

Polyploidy is thought to be a mechanism by which plants can adapt to stressful conditions, providing increased genetic variation and a better chance to evolve beneficial adaptations (Van de Peer et al., 2021, 2017). Amphicarpic species are reported to grow in disturbed habitats (Cheplick, 1987), raising the possibility that polyploidy might have contributed to the *de novo* evolution of amphicarpy in *C. chenopodiifolia* as an adaptation to this stressful environment. Alternatively, geocarpy might have been a trait inherited from a progenitor species, contributing to the evolution of amphicarpy as a bet-hedging strategy for seed dispersal in variable conditions. Although geocarpy has not been reported in other Cardamine species, this trait is present in the distantly related species *Geococcus pusillus* J. Drum. in the Brassicaceae family (Cheplick, 1987). Whether a very wide cross, or an as yet undescribed geocarpic species in Cardamine, contributed to the origin of *C. chenopodiifolia*, are questions to be addressed in future studies. In this regard, the *C. chenopodiifolia* genome is a valuable genomic resource to study the evolution and development of amphicarpy.

## Acknowledgments

We thank R. Mercier and A. Kalde for help performing chromosome spreads, B. Huettel for PacBio sequencing and W. Faigl for technical assistance. This work was supported by Swiss National Science Foundation fellowship P500PB_203021 to A.E. A.H. gratefully acknowledges support from a Max Planck Society core grant to the Department of Comparative Development and Genetics

## Author contributions

Conceptualization, A.E., M.A. and A.H.; Investigation, A.E., M.A., N.T., M.V. and M.P.A.; Writing, A.E. and A.H.; Funding Acquisition, A.E. and A.H.; Supervision, P.N., X.G. and A.H.

