## Supplemental Figures S1 and S2 for "Polyploid genome assembly of *Cardamine chenopodiifolia*"

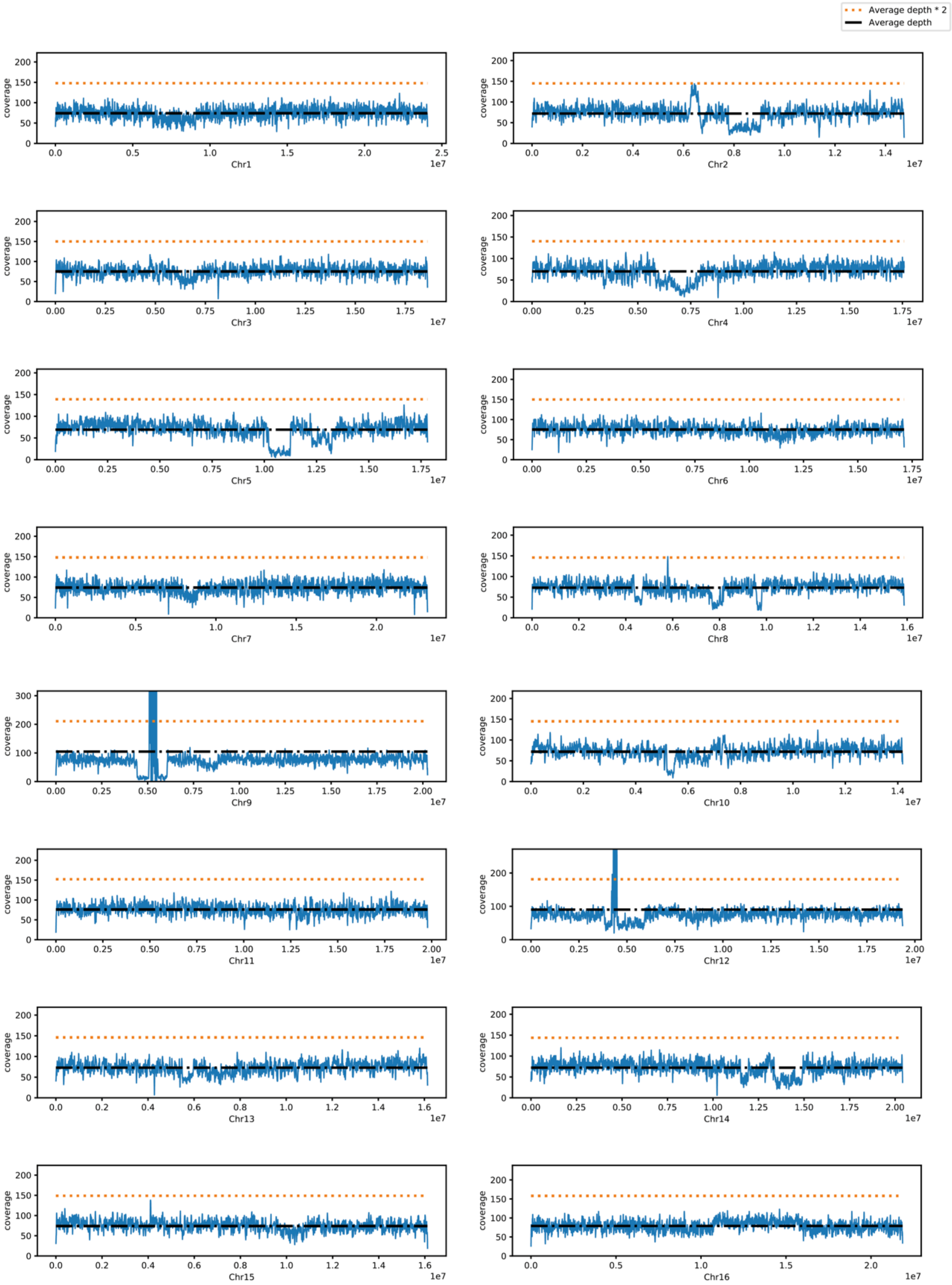


**Figure S1: Depth of coverage for chromosome 1 to 16**

Blue lines indicate the depth of coverage along the 16 first chromosomes calculated with minimap2. Dotted black lines represent the average depth, while dotted orange lines represent twice the average depth.

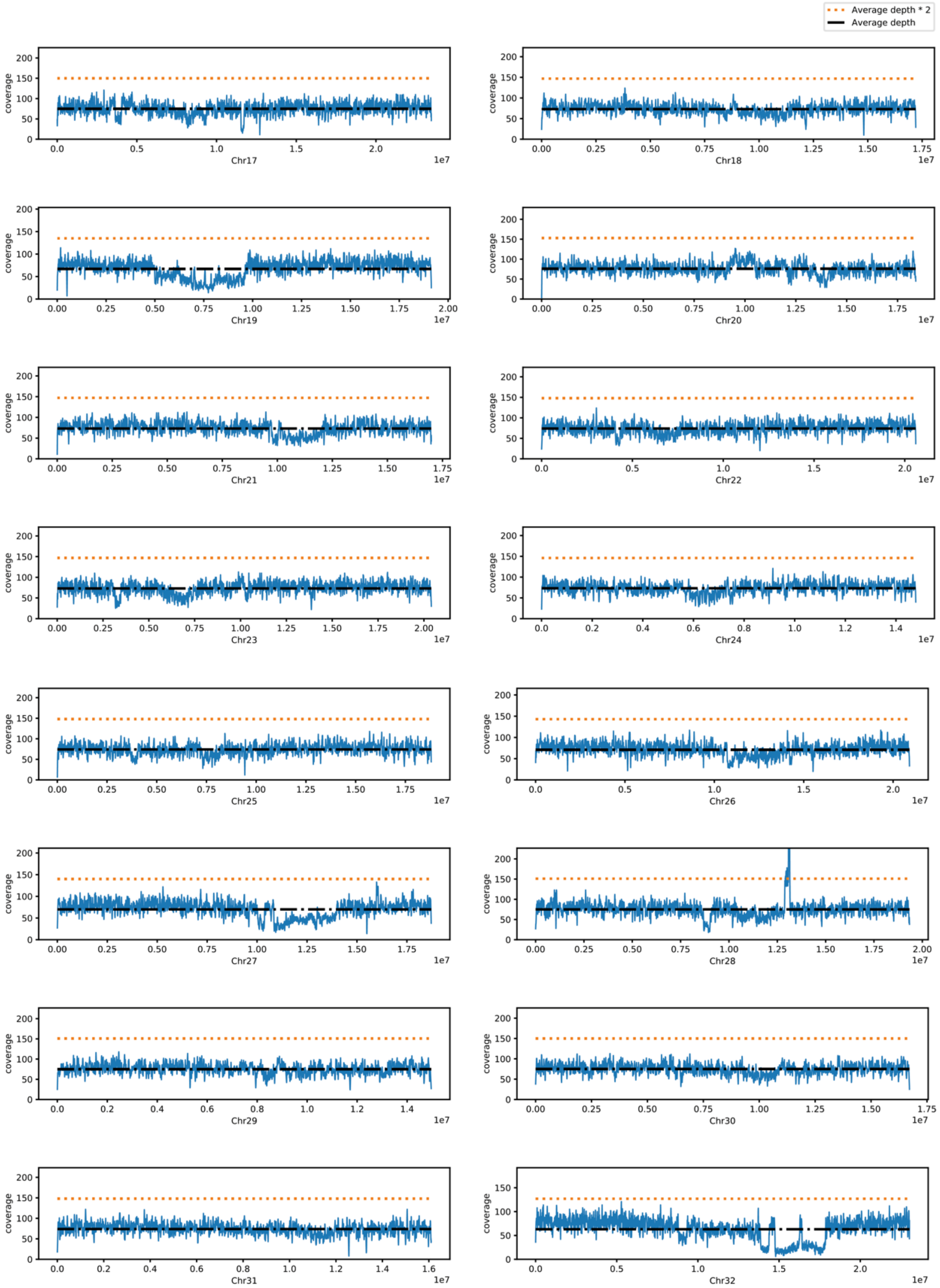


**Figure S2: Depth of coverage for chromosome 17 to 32**

Blue lines indicate the depth of coverage along the 16 last chromosomes calculated with minimap2. Dotted black lines represent the average depth, while dotted orange lines represent twice the average depth.
